# ezscore-f: A Set of Freely Available, Validated Sleep Stage Classifiers for Forehead EEG

**DOI:** 10.1101/2025.06.02.657451

**Authors:** William G. Coon, Paul Zerr, Griffin Milsap, Niloy Sikder, Michael Smith, Martin Dresler, Matthew Reid

## Abstract

The increasing availability of wearable forehead EEG devices, such as the Hypnodyne ZMax, open-source DCM, CGX PatchEEG, and many others, has significantly expanded opportunities for convenient, at-home sleep monitoring. However, most publicly available sleep classifiers are trained on scalp EEG from traditional polysomnography (PSG) data, and thus generalize poorly to forehead EEG due to differences in electrode placement, referencing montages, and higher susceptibility to artifacts from user movements and electrode displacement. Conventional classifiers typically do not explicitly account for these artifacts, resulting in inaccurate and misleading sleep stage scoring. To address this gap, we developed a suite of artifact-aware sleep stage classifiers trained specifically using forehead EEG data, leveraging two comprehensive datasets—Wearanize+ and Donders2022—that contain concurrent forehead EEG and clinical PSG recordings. We further introduce two classifier variants: one optimized for real-time applications that operates directly on raw EEG amplitudes, and another optimized for offline analysis utilizing normalized EEG signals. Validation results indicate robust and reliable classification performance across standard sleep stages (Wake, N1, N2, N3, REM), along with effective identification of artifact epochs. Importantly, the developed classifiers generalize well to forehead EEG devices beyond the original training platform. These validated classifiers are freely available to the sleep research community through the open-source ezscore-f package, providing versatile and practical tools for forehead EEG-based sleep stage analysis.

## I. INTRODUCTION

The rapid proliferation of compact and portable EEG wearables, particularly forehead-mounted devices like the Hypnodyne ZMax, open-source DCM, CGX PatchEEG, and many others, has significantly expanded possibilities for convenient, longitudinal, and at-home sleep monitoring. Compared to traditional laboratory-based polysomnography (PSG), these devices offer non-invasive, cost-effective, userfriendly alternatives suitable for large-scale deployment, which in turn has the potential to greatly accelerate the field of sleep science by providing a way to collect meaningful sample sizes in more ecologically relevant settings. Recent studies validate the capability of such devices to capture sleep EEG dynamics effectively for both macro- and microarchitectural analyses, and concurrently, automated sleep staging approaches using forehead EEG have demonstrated promising results that encompass both proprietary and opensource methods [1]–[4].

Despite these advances, important challenges remain. Due to the highly skilled and labor intensive nature of manual sleep staging, automated sleep stage classifiers are particularly important for large-scale at-home studies. However, most high-performing publicly available sleep classifiers are trained exclusively on standard PSG data [5]–[16], which differ markedly from forehead EEG data in their signal characteristics, referencing montage, and susceptibility to artifact. Consequently, these models frequently underperform when applied to forehead EEG. Recent forehead EEG-specific classification strategies have demonstrated promising results [17], often by decoupling classifier design from traditional sleep staging conventions [3], [4]. However, such approaches depart significantly from standard scoring guidelines (Wake, N1, N2, N3, REM), limiting their comparability with established PSG-based sleep metrics and the broader sleep literature. Moreover, many of these techniques use rule-based decision trees to classify sleep stages, which does not capitalize on the substantial advancements in deep learning that have emerged in recent years.

Perhaps the most crucial obstacle to overcome with forehead EEG is the substantial presence of signal artifacts inherent to wearable devices. Most existing sleep stage classifiers implicitly assume clean, artifact-free EEG—a problematic assumption in real-world settings where wearable EEG often suffers from movement-induced noise, intermittent signal dropout, and electrode displacement. Unlike multi-channel PSG systems that offer redundancy across multiple electrodes, forehead EEG devices typically depend on just two channel derivations, providing minimal tolerance to signal corruption. As a result, without explicit artifact handling, classifiers routinely continue (inappropriately) to assign sleep stage labels to unusable data segments, producing misleading hypnograms and distorting derived sleep statistics.

In response, this manuscript introduces `ezscore-f`, a suite of artifact-aware^1^ neural network classifiers explicitly optimized for sleep staging using forehead EEG data. Leveraging two large-scale datasets for training, Wearanize+ [18] and Donders2022 [1], `ezscore-f` models distinguish the five canonical sleep stages (Wake, N1, N2, N3, REM) and explicitly identify epochs dominated by signal artifacts. This artifact-aware classification approach significantly enhances the reliability and interpretability of automatically generated hypnograms, ensuring derived sleep metrics accurately reflect only valid EEG segments.

To address diverse research and clinical needs, we also provide two model variants: one optimized for real-time classification using raw EEG signals (presented to the classifier in microvolts), and another designed for offline batch analysis using normalized EEG inputs. Initial cross-device validation highlights robust generalization, demonstrating that models trained on the Hypnodyne ZMax can effectively generalize to recordings from other forehead EEG devices without additional classifier training (fine-tuning). All developed classifiers, preprocessing pipelines, and example usage scripts are openly accessible through the ezscore-f package, encouraging widespread adoption and collaborative refinement within the sleep research community.

## II. METHODS

### A. Data Acquisition and Labeling

Data in the Wearanize+ and Donders2022 datasets were collected using Hypnodyne ZMax forehead EEG devices (Hypnodyne Corp., Sofia, Bulgaria). Participants attached the device to the forehead using an adhesive electrode gel sticker incorporating all four EEG sensor contacts (Fig. 1A). The ZMax device’s electrode configuration (see Fig. 1) uses four forehead electrodes that, after differential referencing, reduce to two EEG derivations (Left and Right). Signals from both derivations were recorded overnight at a sampling frequency of 256 Hz.

**Fig. 1.**
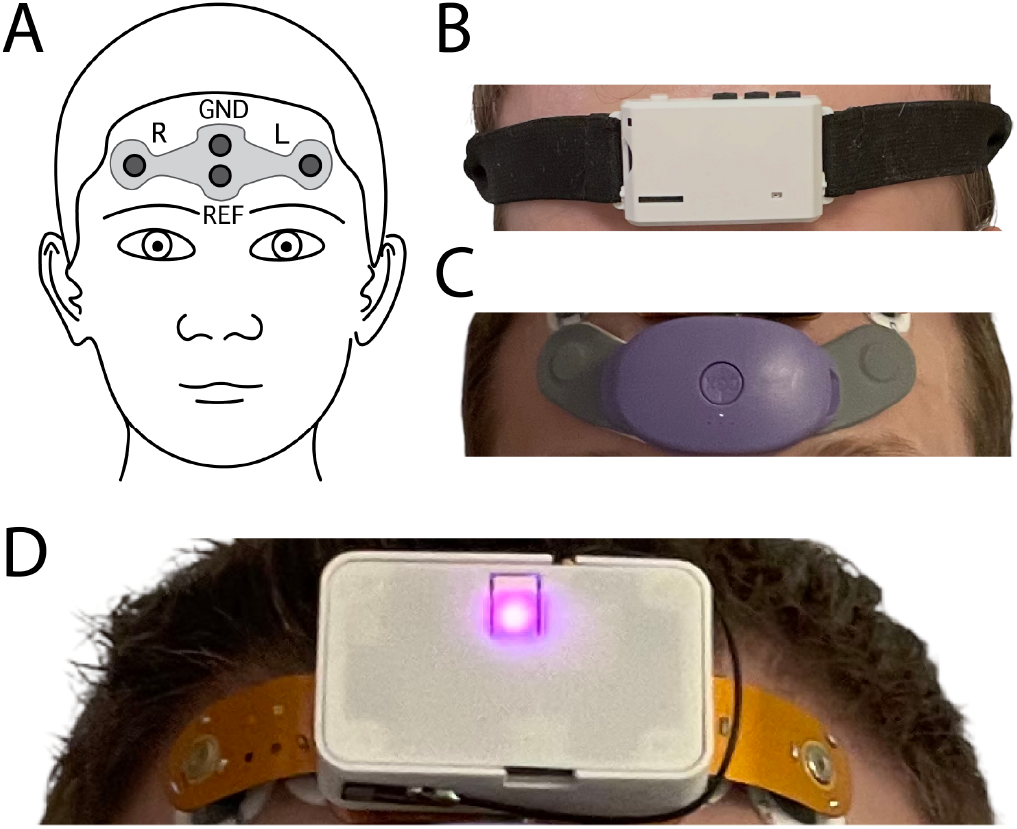
Electrode sensor locations in forehead EEG devices. **A**. Sensor layout common to many forehead EEG devices such as the Hypnodyne Zmax shown in **B. C**. PatchEEG by CGX. **D**. DCM (JHU/APL), an opensource forehead EEG patch designed for realtime processing and sleep research [19].

Each PSG recording was manually scored by an expert rater following standard American Academy of Sleep Medicine (AASM) guidelines [20]. Sleep was annotated in standard 30-second epochs into five classes: Wake, N1, N2, N3, and REM.

### B. Preprocessing and Artifact Labeling

Prior to analysis, we minimally preprocessed data with a high pass filter at .5Hz and downsampled to 64Hz. To explicitly identify epochs severely impacted by artifacts, we computed epoch-wise EEG signal quality metrics, including mean amplitude, median amplitude, and Hjorth parameters (Complexity and Mobility). Thresholds for artifact identification were determined empirically, informed by expert scorer assessment and validated against manual inspection. Specifically, epochs with mean amplitudes below 1.4 µV (indicative of poor electrode contact) or above 200 µV (associated with high-amplitude artifacts such as movement) were marked as artifact epochs (Fig. 2A). These thresholds were uniformly applied across all data.

**Fig. 2.**
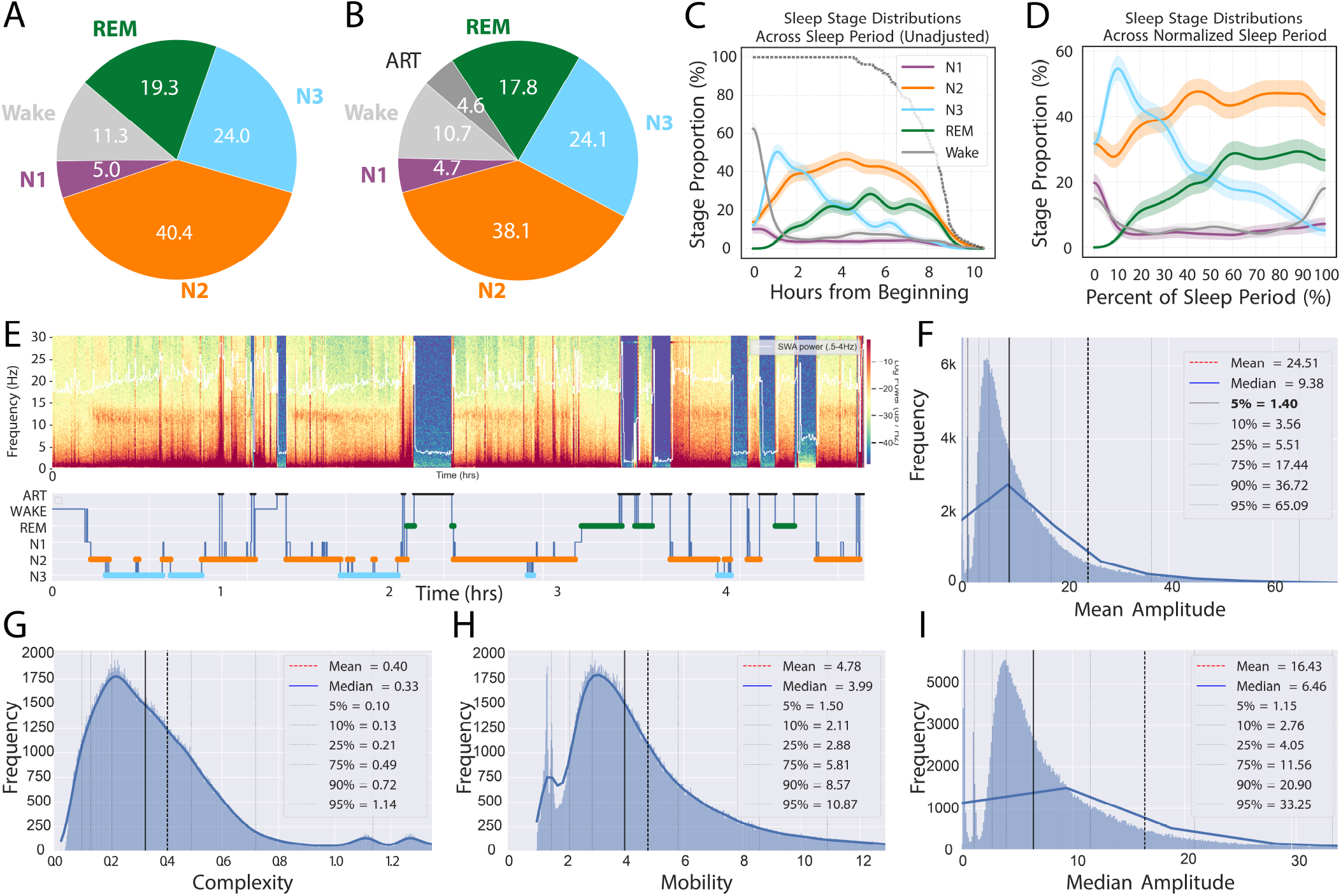
Sleep and artifact statistics. **A**. PSG-derived sleep stages in the dataset (percent total). **B**. Same source data as in (A), but with epochs identified as artifact labeled accordingly (see Methods). **C**. Sleep stage densities across the night. Lines show mean*±* **SEM** across all participants (n=160) **D**. Timenormalized densities, with time zero defined by the first epoch of sleep in each record. **E**. Example of visibly apparent artifact automatically identified by the artifact detector. At top, a spectrogram reveals intermittent signal loss / low-amplitude noise, likely due to poor electrode contact. At bottom, the PSG-derived hypnogram, with artifact periods marked as a 6th class (see Methods) for the artifact-aware variants of `ezscore-f` to learn. **F**. Distribution of mean absolute amplitude in each 30-second epoch in the data. Artifact detection was thresholded at less than the 5th percentile (1.4µV) or greater than 200µV. **G**. Hjorth complexity distribution. **H**. Hjorth mobility distribution. **I**. Median amplitude distribution.

### C. Training and Validation data

This study analyzed two publicly available datasets, Wearanize+ and Donders2022, which together consist of160 overnight recordings of forehead EEG acquired with concurrent polysomnography (PSG). The Wearanize+ dataset included single recordings per participant (*n* = 96 individuals, *m* = 96 recordings), while Donders2022 consisted of multiple recordings per participant (1 to 3 per participant) collected one week apart (*n* = 30 individuals, *m* = 64 recordings). The total number of participants was *n* = 126 individuals. Participants in both studies provided informed consent and the studies were approved by the institutional research ethics committee, METC Oost-Nederland (2014/288), in accordance with DCCN blanket approval, protocol ‘Imaging Human Cognition’ (NL45659.091.14).

Separate analyses were conducted for a 5-class sleep stage classifier (Wake, N1, N2, N3, REM; definitions based on AASM criteria [21]), and a 6-class classifier that explicitly incorporated an additional artifact (ART) class. A summary of sleep stage distributions in the data is provided in Figure 2.

To establish an upper-bound reference performance level for forehead EEG classification using 5-stage, artifact-naive classifiers, we separately trained an additional 5-class model variant using a subset of artifact-free records (>99% artifact-free, epoch-wise, visually confirmed by expert rater). This allowed assessment of the baseline (artifact-naive) model’s performance under ideal conditions, which could then be compared against the performance of the same model trained on data of realistic, real-world quality (i.e., data which include unusable signal segments). This also allowed comparison against the artifact-aware models trained and tested on the exact same real-world quality training dataset.

### Model Training

We adapted a hybrid neural network architecture combining convolutional neural networks (CNNs), transformer encoder layers, and recurrent neural networks (RNNs; specifically bidirectional long short-term memory [LSTM] networks). This design leveraged the CNN layers for spatiotemporal and frequency-domain feature extraction, transformer layers to capture fine-grained temporal dependencies within epochs, and RNN layers to model temporal dynamics across epochs (Fig. 3).

**Fig. 3.**
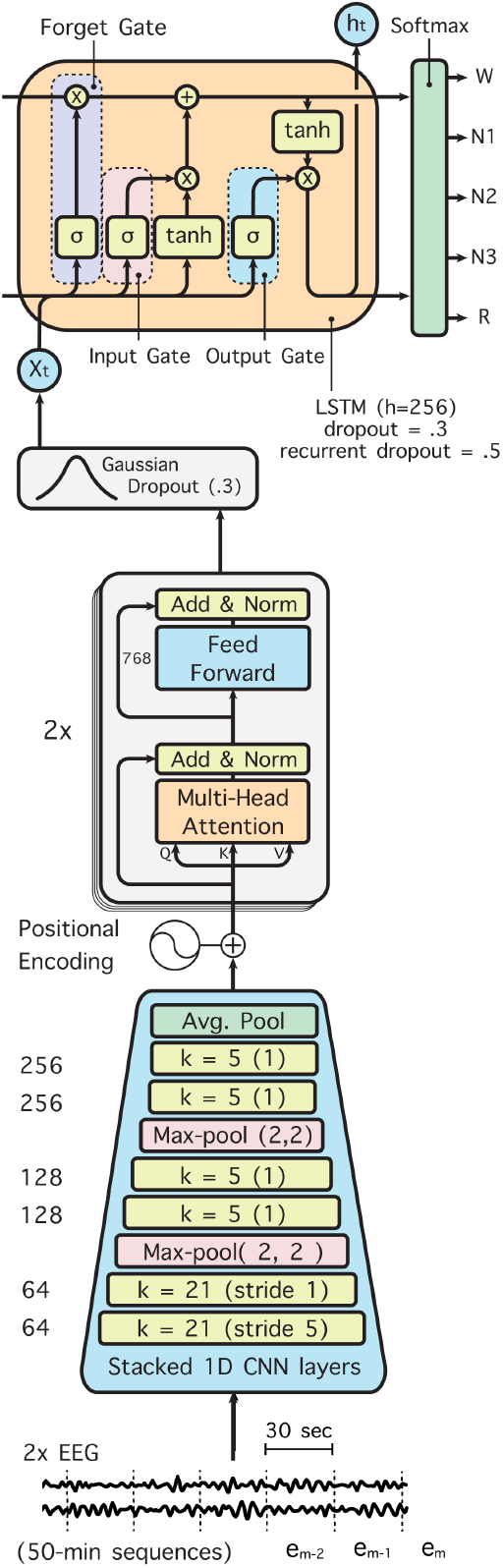
Architecture of the hybrid CNN-transformer-RNN neural network used in both 5and 6-class models. Feature extraction is performed by a CNN front end, passing high-dimensional representations to a transformer stack to encode temporal dependencies across data samples in each epoch. The transformer-encoded features are then passed to a bidirectional RNN, implemented as a LSTM, to capture temporal dependendies across epochs within each 50-minute input segment. The model outputs a sleep stage label for each 30-s epoch in the 50-minute (100epoch) sequence (5-class model shown; 6-class model adds an “ART” class output from the penultimae softmax layer.

One innovation in the classifier architecture was the parallelization of the transformer stack so that each parallel transformer, receiving the same output from the CNN feature extractor, could focus on different types of temporal dependencies in those features. One transformer stack terminated in a max-pooling layer, which emphasized brief signal transients like K-complexes and sleep spindles, while the other terminated in a global average pooling layer, which better attended to steady-state trends in data such as contiguous slow waves during N2 or N3 sleep. Both views were concatenated and then submitted to the RNN for across-epoch sequence modeling.

Given this architecture and the hyperparameters used in this study, the model thus views up to 50 minutes of sleep at a time, assigning sleep stage labels in sequence while considering patterns across the entire 50-minute view. Importantly, during inferencing, the model can operate on as few as a single 30-second epoch at a time. This dynamically scalable view length is especially advantageous for real-time scenarios.

Models were implemented in Python 3.10 using TensorFlow 2.13.0, and Keras-NLP 0.9.3. Classifier training involved minimizing categorical cross-entropy loss through stochastic gradient descent optimization (Adam optimizer). Training proceeded for 100 epochs, with an initial learning rate of 0.001 reduced to 0.0001 after eight epochs. Categorical cross-entropy (*H*(*P*)) between labeled classes was minimized as:

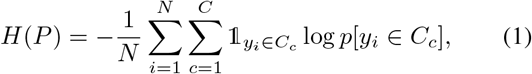

with entropy *P*, number of examples *N*, number of classes *C* (here, *C* = 5 or *C* = 6 sleep/wake labels), and *y*_*i*_ a single example (30-second epoch).

Unnormalized data (e.g., raw µV values) can be advantageous for real-time processing because standard normalization techniques often require access to the full overnight recording to compute measures of central tendency and variability—a requirement incompatible with real-time applications. However, normalization of time series data typically enhances classifier performance and is therefore preferred in retrospective analyses, where the complete dataset is available and real-time constraints do not apply. Accordingly, we developed two variants of the artifact-aware classifier: one operating directly on raw EEG amplitudes (in microvolts, µV) optimized for real-time use, and another using medianinterquartile-range (IQR)-normalized EEG signals clipped at ±20 times the IQR (as in [5]), optimized for offline analyses. Models were trained using a 90/10 train/test set distribution, in which 90% of the total dataset was used for classifier training and the remaining 10% were used to evaluate model performance.

### E. DCM data

To assess generalization to EEG devices beyond the Hypnodyne ZMax, we collected two overnight recordings of co-acquired PSG and a comparable forehead EEG device (DCM [19], Johns Hopkins University Applied Physics Laboratory, Laurel, MD; shown in Fig. 1; participant provided informed consent to participate in this study, approved by the Johns Hopkins University’s Institutional Review Board). DCM data were digitized at 250 Hz using custom software built upon the open-source ezmsg platform [22], designed explicitly for low-latency real-time EEG processing. Identical preprocessing steps (resampling, filtering, and normalization) used for the ZMax data were applied to DCM recordings. PSG data from these overnights were scored by an RPSGT and reviewed by a licensed sleep medicine physician at the Johns Hopkins Centre for Interdisciplinary Sleep Research and Education.

## III. RESULTS

### A. Artifact-Naive Model (Baseline)

When trained on the full forehead EEG dataset (including artifacts), the baseline artifact-naive classifier showed relatively poor overall accuracy (63.9%, Cohen’s *κ* = 0.49; Fig. 4A, Table I). Notably, accuracy was substantially compromised for Wake (56.0%), N1 (7.0%), and REM (24.2%). The model frequently misclassified REM epochs as either Wake or N2 sleep, underscoring the difficulty posed by forehead EEG artifacts on traditional classification approaches. Classification of deeper sleep stages, N2 and N3, remained robust (77.9% and 84.9%, respectively).

**Fig. 4.**
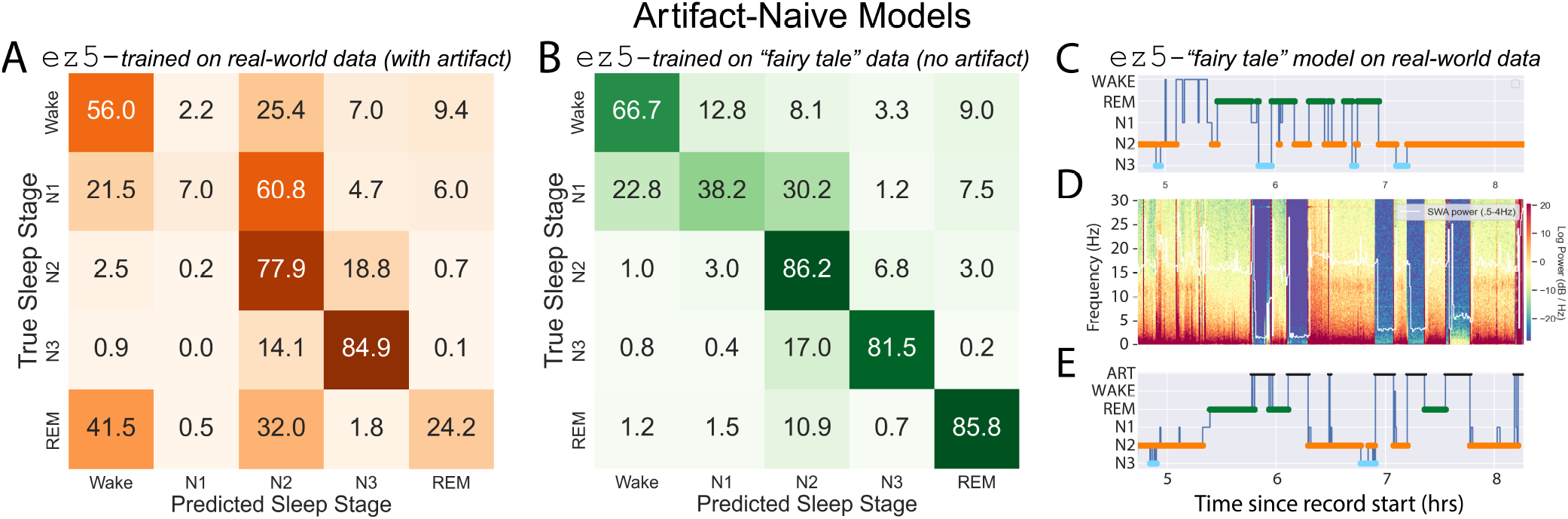
Artifact-heavy data pose classification challenges: **A**. Accuracy (%) from 5-class model, including dirty data. The same train/test sets were used, but artifact remained unidentified in the training labels, injecting substantial noise into the training set and devastating the model’s capacity to distinguish Wake and REM. **B**. Accuracy from 5-class model, excluding dirty data. In this example, dirty records were removed from the training set. Superficially this improves performance, but leads to spurious results when applied to noisy data. **C**. Single example using the classifier in (B). Note how the confusion matrix in (B), which was derived from all 19 participants in the held out test set, looks superb at first glance. The classifier continues to report sleep stages despite there being no meaningful information in signal dropout regions, while also substantially distorting predicted sleep stages in viable signal regions surrounding areas of artifact. This issue may go unnoticed in large datasets, potentially skewing population statistics significantly.) **D**. EEG spectrogram. **E**. Ground truth sleep stages from PSG and artifact labeling procedure (see Methods).

**TABLE 1.** MODEL VARIANT SUMMARY.

| Model | Artifact Aware? | Trained on | Tested on | Purpose | $\kappa$ |
| --- | --- | --- | --- | --- | --- |
| ez5 | art-naive | art-free | art-heavy | offline | .49 |
| ez5ft | art-naive | art-free | art-free | offline | .73 |
| ez6rt | art-aware | art-heavy | art-heavy | real-time | .71 |
| ez6 | art-aware | art-heavy | art-heavy | offline | .74 |

### B. Artifact-Naive Model (“Fairy Tale” Data)

To establish best-case performance benchmarks, we trained an additional artifact-naive model on artifact-free (“fairy tale”) data only. This substantially improved overall performance (on artifact-free data), achieving 80.6% accuracy and a Cohen’s *κ* of 0.73 (Fig.4B). Performance markedly improved for Wake (66.7%), N2 (86.2%), N3 (81.5%), and REM (85.8%). However, N1 accuracy remained relatively low (38.2%). Importantly, this artifact-naive model, when presented with unseen data containing substantial artifact contamination, continued to assign sleep stages despite the absence of meaningful EEG signal, resulting in misleading hypnograms (Fig.4C-E).

### C. Real-Time Artifact-Aware Model (µV Input)

To explicitly handle artifacts, we trained an artifact-aware classifier that incorporated a sixth class for artifact identification, and operates directly on raw EEG amplitudes (suitable for real-time applications). This model significantly improved classification accuracy relative to the baseline artifact-naive model, achieving an overall accuracy of 79.2% (Cohen’s *κ* = 0.71; Fig. 5A) on the 6-class problem. Accuracy notably increased for REM (91.3%), N2 (82.7%), and N3 (82.7%), though Wake accuracy (56.8%) and artifact classification accuracy (35.0%) remained relatively modest. N1 accuracy improved slightly (33.6%), but remained limited due to class imbalance.

**Fig. 5.**
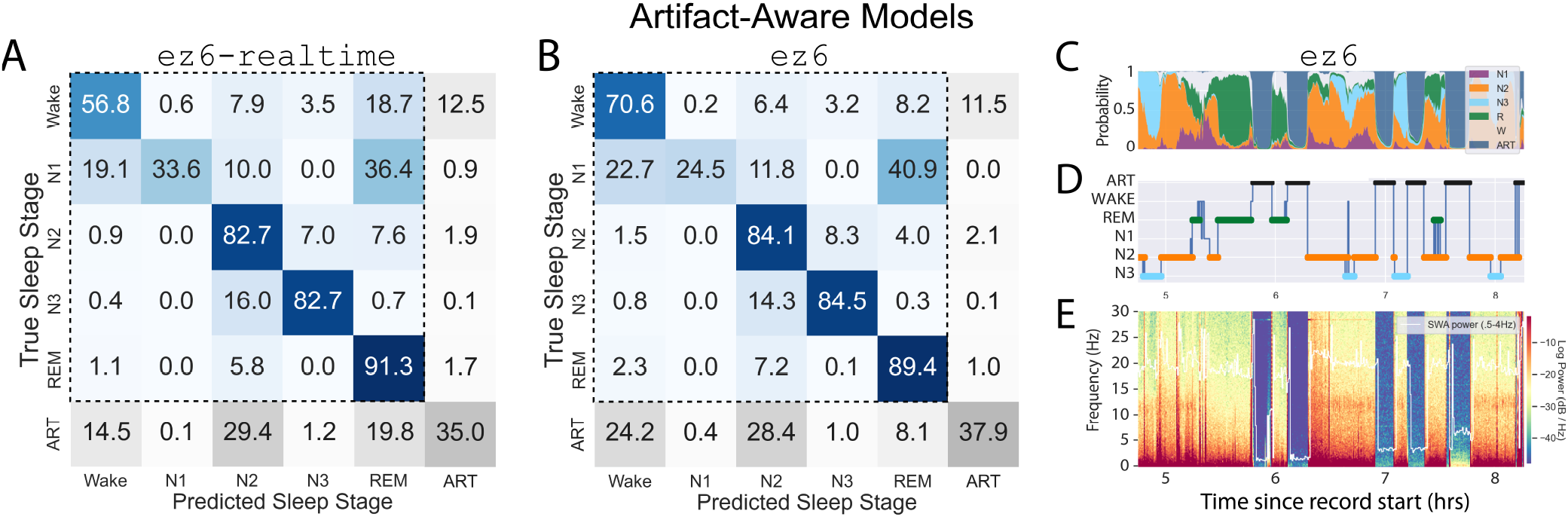
Adding a 6th class rescues performance and avoids assigning sleep stages to periods rendered unusable by artifact: **A**. Accuracy (%) from the 6-class model. This model is optimized for real-time processing, accepting input in units of raw microvolts (µV). **B**. Accuracy from the 6-class model trained using median/IQR-normalized signals as input (best for offline classification). This further improves performance. **C**. Classifier output probabilities (“hypnodensity”). Note confident identification of artifact. **D**. Classifier output hypnogram. Artifact is identified as an independent class, allowing summary sleep statistics to be calculated without any skew injected from artifact-driven mis-scores. **E**. EEG spectrogram exhibiting clear periods of intermittent signal loss.

**Fig. 6.**
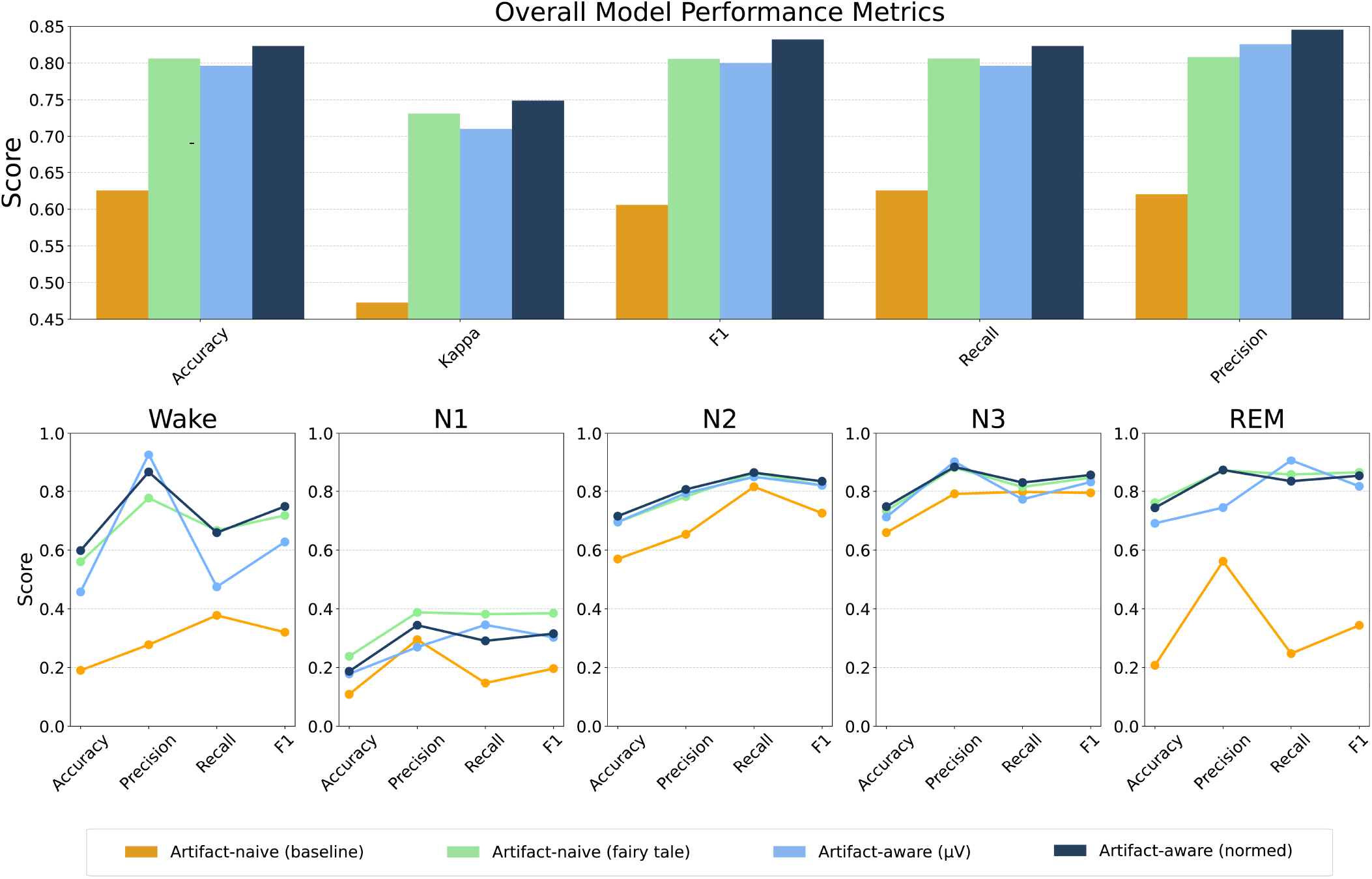
Model performance comparison. *Top:* Overall performance on five measures of model performance: Accuracy, Cohen’s kappa (*κ*), F1-score, Recall, and Precision. Note how the 5-class model trained on largely artifact-free (“fairy tale”) data deceptively appears to performs well (light green bar). If applied to data with high levels of artifact contamination, spurious sleep stage labels will be assigned to those portions (see Fig. 4 for example). Adding a sixth class for artifact rescues performance and adds the desirable property of being valid to apply to real-world data, even when contaminated by long periods of signal artifact. *Bottom:* Metrics by sleep stage.

It is important to consider that, when comparing performance scores across classifiers posed with different numbers of classes to learn, chance probability changes with each change in the number of classes. Hence direct comparison of e.g., accuracy scores can be misleading when a different number of classes was learned by one compared to the other. Considering performance only within the five sleep stage labels puts both 6- and 5-class models on equal footing, making the comparison fair and more readily interpretable. Overall accuracy for the artifact-aware model on the 5-class sleep stage problem was 80.3% (*κ* = 0.73).

### D. Offline Artifact-Aware Model (Normalized Input)

The offline version of the artifact-aware classifier operated on end-to-end normalized EEG signals, which further improved performance, achieving an overall accuracy of 81.3% (Cohen’s *κ* = 0.74; Fig. 5B). Wake accuracy substantially increased to 70.6%, and accuracies for N2 (84.1%), N3 (84.5%), and REM (89.4%) were robust. Artifact accuracy slightly improved to 37.9%, while N1 accuracy decreased to 24.5%. Thus, normalization clearly benefited Wake stage detection and marginally reduced REM accuracy compared to the real-time variant. Still, REM accuracy remained high overall. Overall performance for the 5-class sleep stage problem was the highest we observed in this study, at 82.3% (*κ* = .75; Fig. 5).

### E. Cross-Device Generalization

The `ez6` artifact-aware classifier, when applied without modification or fine-tuning to DCM EEG data, exhibited an overall accuracy of 76.4± 0.2%, with a mean *κ* = .65± .02. One overnight is shown in Fig. 9.

### Balancing Performance Gains Across Classes

While the overall performance of the artifact-aware classifiers was comparable to the best available state-of-the-art polysomnography (PSG) classifiers [5], their performance on minority sleep stage classes, such as N1 and ART, was for Interdisciplinary Sleep Research`ez6` model using a hybrid loss function. The loss function combined two objectives: standard cross-entropy loss, accounting for 90% of the total, and a “soft” Cohen’s *κ* approximation (“soft *κ*”) constituting the remaining 10%. Cohen’s *κ* is a widely-used metric for inter-rater agreement in sleep research because it adjusts agreement measures for chance. We chose the 90/10 weighting ratio strategically: introducing the “soft *κ*” term at a moderate level improved performance on minority classes without destabilizing training, which occasionally occurred when this component was weighted more heavily. The use of “soft *κ*”—employing probability densities (softmax outputs) rather than discrete class predictions (argmax)—facilitates gradient-based optimization during model training. The final loss function can be written as:

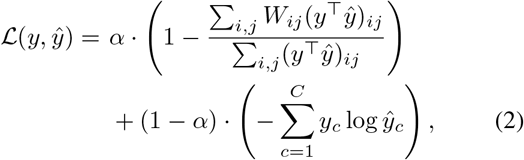

where

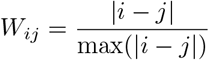

with *y* and *ŷ* representing the one-hot encoded ground truth labels and the predicted class probabilities, respectively, for a sleep stage classification task with *C* total classes. The first term in the loss is a “soft” differentiable approximation of the weighted Cohen’s kappa metric, controlled by the hyperparameter *α* ∈[0, 1]. The weighting matrix *W*_*ij*_ penalizes disagreements between classes *i* and *j* proportionally to their ordinal distance. The second term is the standard categorical cross-entropy loss.

The hybrid loss function substantially improved model performance (Fig. 7), with a final overall (6-class) accuracy of 81.4% (*κ* = .75) and N1 accuracy of 89.6%.

**Fig. 7.**
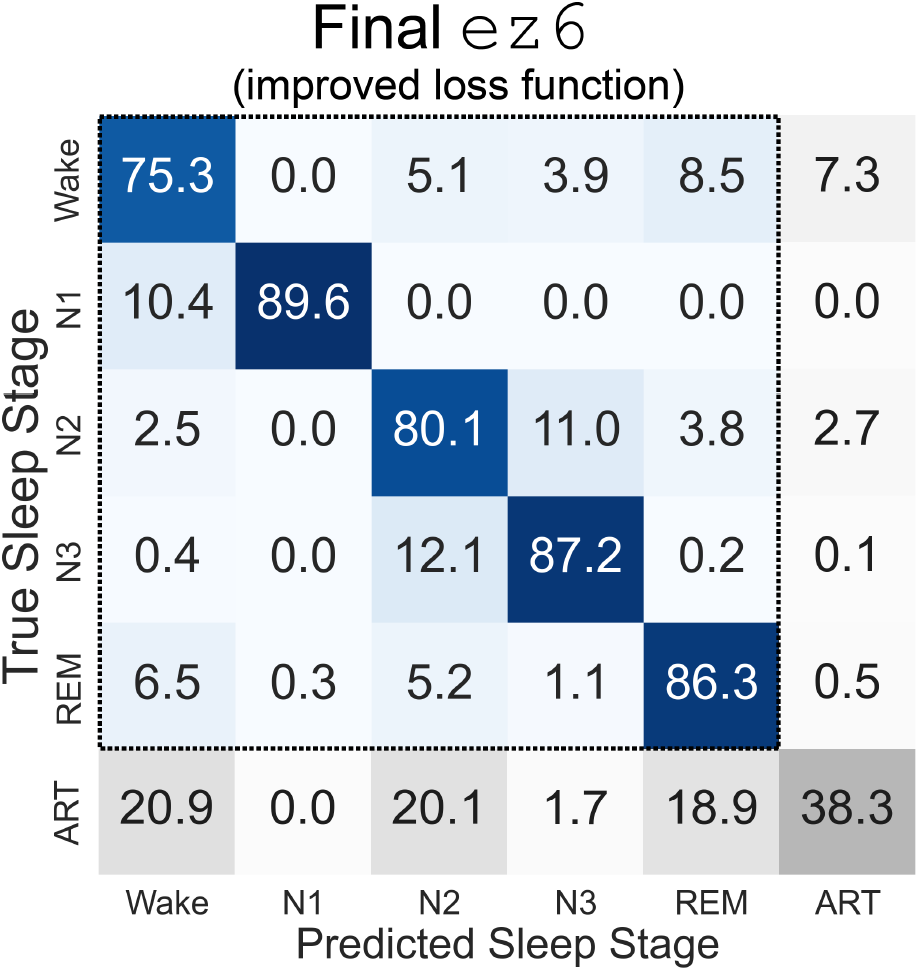
A soft *κ* loss function rescues performance on minority classes. Training the `ez6` model using a hybrid loss function designed to address class imbalance stabilizes performance across all sleep stages.

**Fig. 8.**
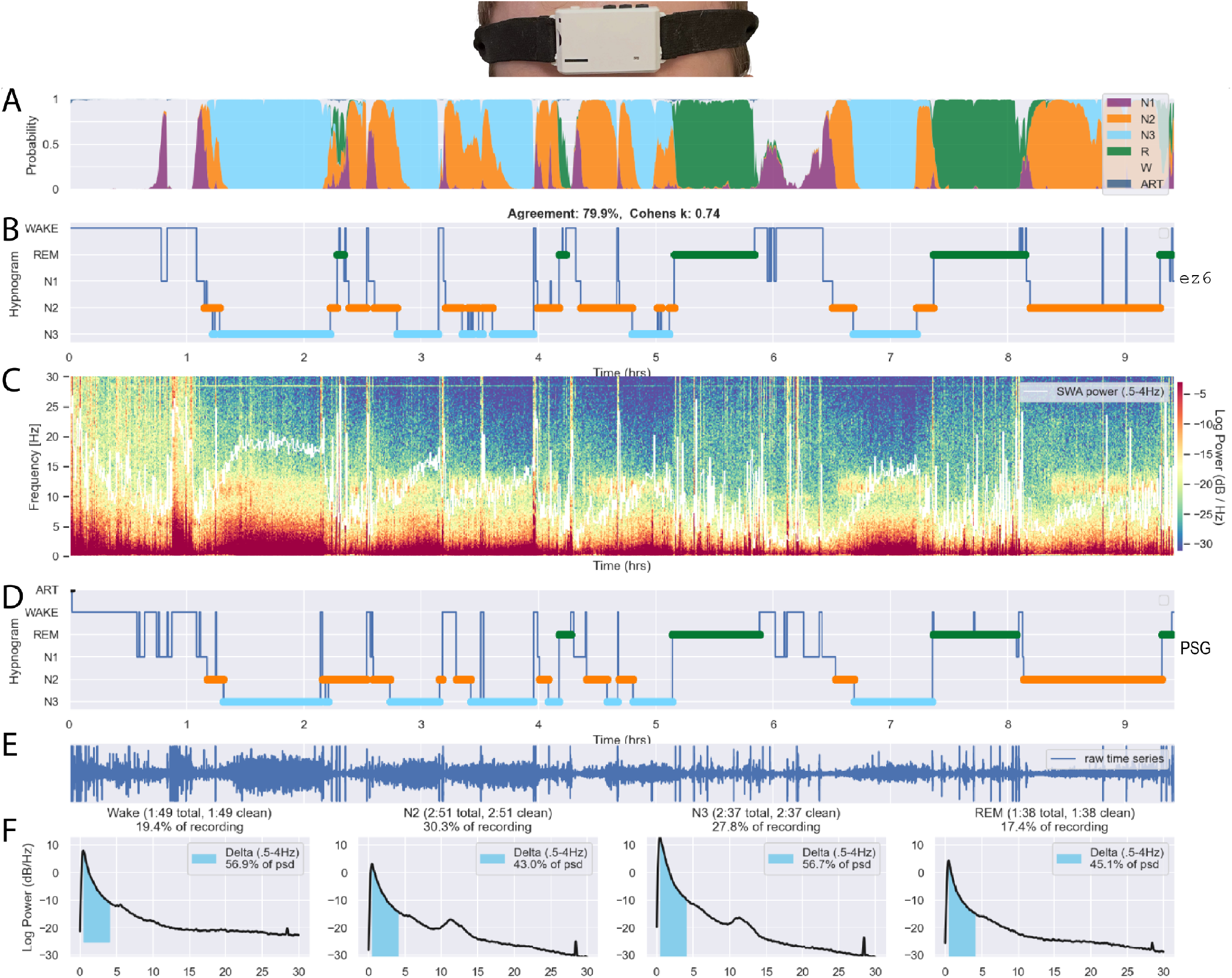
Artifact-aware ez6 model’s validation performance on a single night of ZMax data. Recording time series (E) and spectrogram (C) from the left EEG channel show expected dynamics (ex. decline in slow wave activity (SWA) power over the course of the night (white line in C), signal amplitude increases during slow wave sleep (N3), etc.). Ground truth hypnogram is shown in (D). This is a relatively long and complex record to score, as evidenced by e.g., intermingled Wake and REM (which would be indistinguishable to the human eye using EEG alone), and several arousals correctly scored was Wake and not Artifact. Line spectra in (F) highlight both delta and spindle (sigma; 12-15Hz) band oscillatory bumps in the expected sleep stages (highest delta (here defined as .5-4 Hz; highlighted in turquoise) in N3, biggest spindle/sigma bumps in N2 and N3, broadband power decline and no appreciable oscillatory activity during REM), and also the presence of a narrow-band high-amplitude signal artifact spanning the entire recording. Note that this did not interfere with the model’s ability to correctly identify sleep stages and not mislabel as Artifact.

**Fig. 9.**
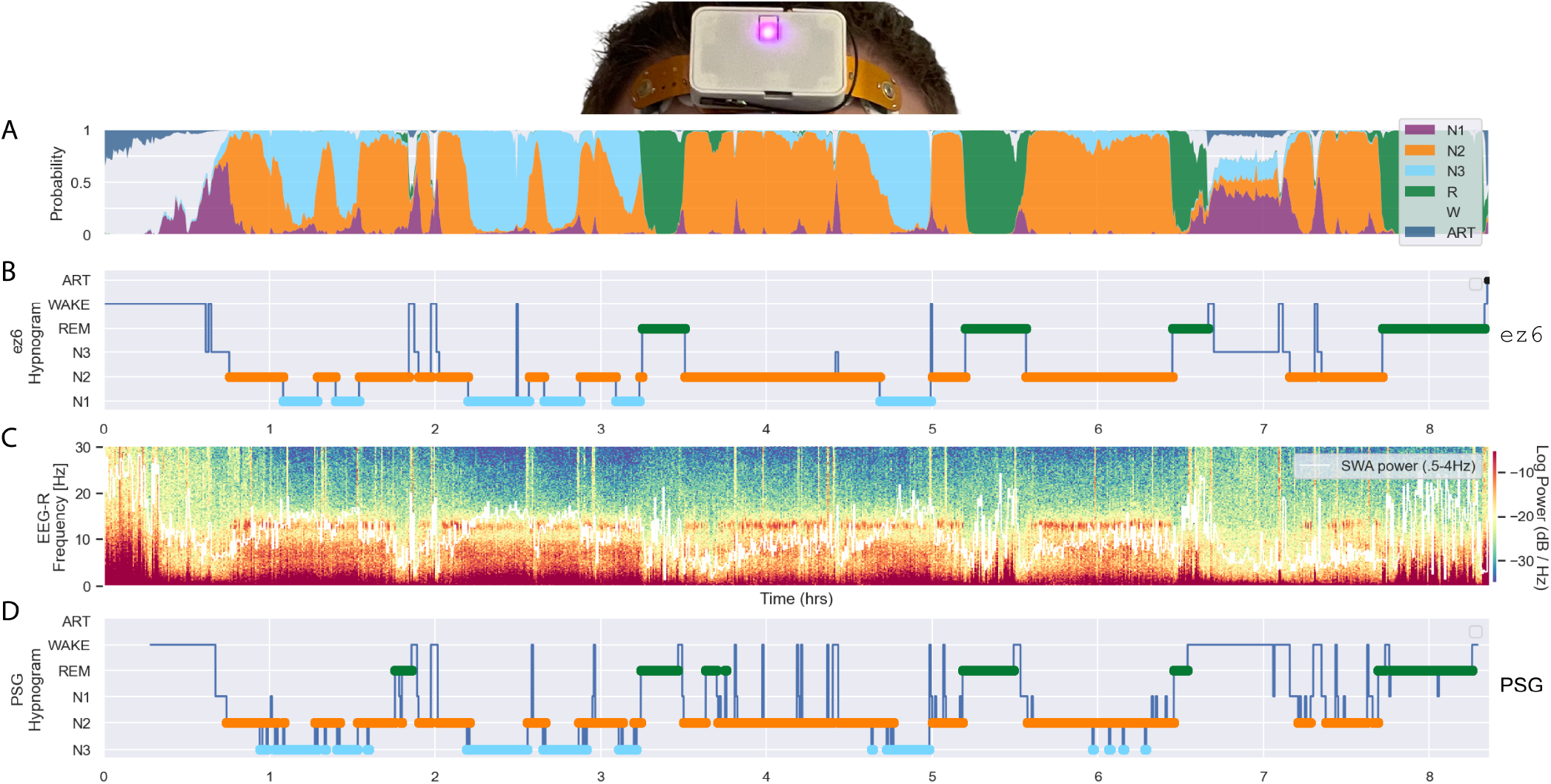
Artifact-aware ez6 model: cross-platform generalization. The `ez6` model demonstrated exemplary performance on a new hardware platform (DCM [19], JHU/APL) despite having been trained on another platform’s data and not having been fine-tuned or otherwise adjusted to adapt to the new platform. **A**. Hypnodensity plot of sleep stage probabilities output by ez6. **B**. `ez6` hypnogram. **C**. DCM EEG spectrogram, representative of typical DCM recording quality. **D**. PSG hypnogram used as ground truth for quantifying performance (in this example, accuracy was 76.6% with a Cohen’s *κ* = .66).

## IV. DISCUSSION

The `ezscore-f` classifiers presented here address key limitations in sleep staging methodologies for forehead EEG devices, delivering robust and validated models designed explicitly for this increasingly popular sleep monitoring approach. Traditional classifiers trained on clinical polysomnography (PSG) data frequently underperform when applied to forehead EEG, primarily due to differences in electrode placement, referencing montages, and heightened susceptibility to artifacts. By explicitly accounting for these artifacts, the proposed classifiers notably enhance the accuracy and reliability of automated sleep staging, effectively eliminating misleading sleep stage assignments during periods of signal degradation or loss.

### A. Impact of Artifact-Aware Sleep Classification

A critical advancement of this approach is the explicit identification and handling of unusable EEG data segments. Conventional classifiers, even those designed specifically for forehead EEG but lacking explicit artifact labeling, typically continue assigning sleep stages irrespective of data quality, thereby risking spurious or fictitious results (see Fig. 4C for on such example). In contrast, the artifact-aware method described herein proactively identifies artifact-dominated epochs, ensuring that sleep staging remains reliable and interpretable, especially in real-world scenarios involving uncontrolled environments. Alternative artifact-handling approaches, such as artifact rejection during pre-processing (for both training and inference data), can provide some explainability advantages over the current approach, as in that case it would be clear which artifact was rejected for what reason. Future work should explore “explainable AI” approaches to further enhance transparency and trust in artifact-aware models. The Shapely Additive Explanations method (SHAP), for example, can explicitly identify decision-driving signal features by using input perturbations and game theory to assign credit to features that most strongly influence model behavior [23]).

### B. Impact of Time Series Normalization

Comparative analyses of real-time and offline variants revealed the significant impact of EEG normalization on classifier performance. Normalization markedly improved accuracy, which underscores the practical trade-offs between accuracy and operational constraints as normalized EEG significantly boosts offline accuracy, whereas real-time applications, such as closed-loop sleep interventions, require classifiers capable of operating directly on raw EEG data. Both classifier variants thus offer distinct advantages tailored to their respective use-cases.

### C. Addressing Class Imbalance in Sleep Stage Classification

One notable limitation identified early in the study was the consistently low classification accuracy for the N1 sleep stage, primarily driven by significant class imbalance (N1 epochs represented less than 5% of total epochs). This was effectively addressed by training models with a custom hybrid loss function sensitive to class imbalance. Future improvements could incorporate oversampling methods or specialized training strategies such as the geometric mean-based loss function proposed by Jones et al. [24]. Such approaches explicitly incentivize the model to accurately classify underrepresented stages, thereby enhancing overall classification robustness and utility.

### D. Enhancing Artifact Identification Accuracy

Artifact classification accuracy, while instrumental in overall performance gains, remained moderate (approximately 35–38%), likely due to treating all artifacts as a single category despite substantial differences in their amplitude and spectral properties. This can be visibly appreciated in lowdimensional projections of high-dimensional class features extracted at the output of the CNN layer of the network (Fig. 10). Introducing separate labels for distinct artifact subtypes (e.g., low-amplitude artifacts due to poor electrode contact versus high-amplitude movement artifacts) may significantly enhance classifier performance. Additionally, future model architectures with increased capacity or complexity could better accommodate this additional classification dimension, further refining artifact detection.

**Fig. 10.**
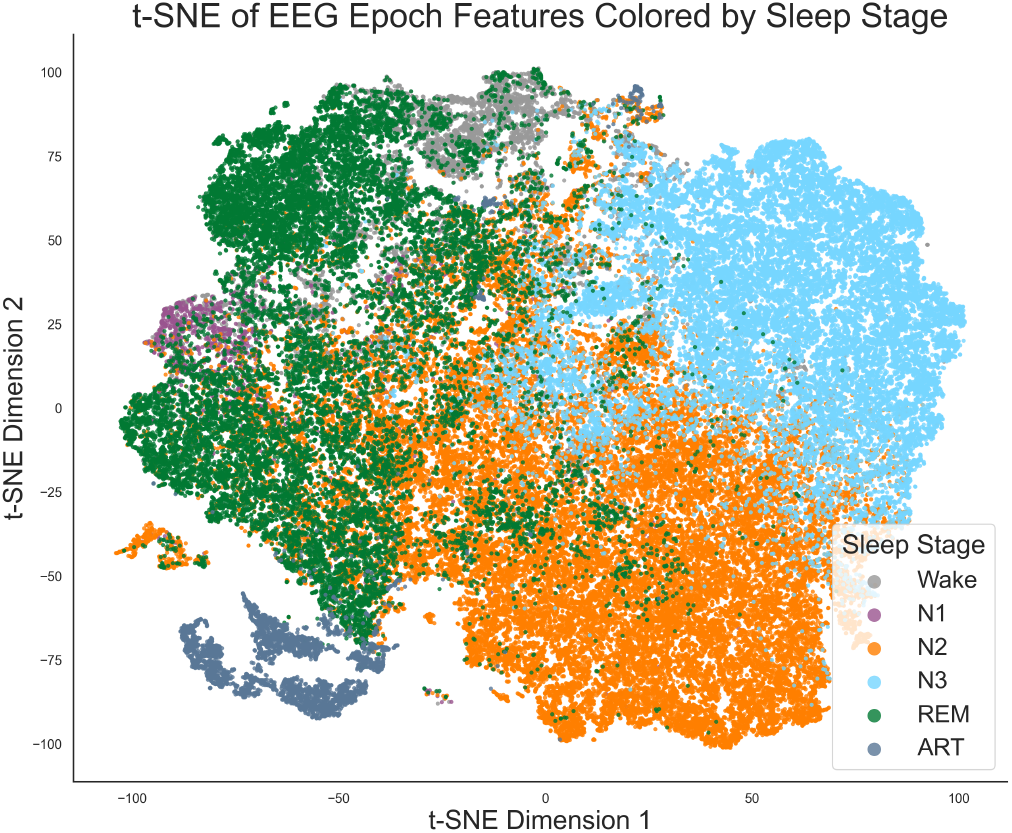
tSNE projection of the high-dimensional representations of sleep stage (class) features, extracted early in the network at the output of the CNN stack. Note that the ART class forms clearly distinct clusters in feature space (lower left), indicative of heterogeneity in the types of artifact represented in the data. Note also that the sleep stages will be substantially more separable at the output of the entire network, after re-encoding by transformers and submission to sequence modeling by the RNN.

### E. Generalization Across Forehead EEG Devices

Classifiers demonstrated preliminary evidence of robust generalization from the Hypnodyne ZMax training platform to another forehead EEG device (DCM), underscoring the potential broad applicability of `ezscore-f`. In this study we were only able to quantitatively test 2 subjects overnight recordings with concurrent DCM and PSG for validation, and further validation in a larger sample is clearly warranted. However the marginal difference in performance (compared to performance on unseen data from the same platform as the training data, i.e., ZMax) suggests that minimal fine-tuning would be needed to achieve optimal results in new hardware, and performance without fine-tuning may already be sufficiently high for many applications. Future work should test this quantitatively, systematically varying the amount of finetuning data needed to achieve a given level of performance on new platforms. Given the growing diversity of forehead EEG devices, further validation across additional platforms and varied recording conditions will be essential. Employing unsupervised domain adaptation and domain generalization techniques (see [25] for review) could further enhance cross-device compatibility, minimizing the need for device-specific retraining and enabling wider adoption of advanced sleep staging capabilities [25], [26].

### F. Freely Available Open Source Release

The classifiers, pre-processing tools, and usage examples provided in the open-source ezscore-f package are intended to facilitate widespread adoption, collaborative improvement, and validation across diverse EEG platforms and research contexts. By making these resources freely accessible, we aim to support advancements in sleep science, facilitate personalized and closed-loop sleep interventions, and promote reliable and accurate sleep monitoring beyond traditional clinical settings.

*To foster early community testing and facilitate iterative improvement, models, fully annotated source code, and tutorials are available at:* **https://github.com/coonwg1/ezscore**

## Footnotes

1 In this context, we use the term “artifact-aware” to refer to models trained on data that includes labeled signal artifacts, enabling the model to recognize and distinguish corrupted segments. In contrast, “artifact-naive” models are trained under the assumption that all input data is artifact-free (whether or not that is actually be the case).

